# Evidence for the ultra-soft brain provided by uniqueness in intrinsic MR elastography

**DOI:** 10.64898/2026.01.23.701281

**Authors:** Marius Burman Ingeberg, Elijah Van Houten, Jaco J.M Zwanenburg

## Abstract

**Introduction:** Brain tissue is exceptionally soft. Recent *in vivo* work, including intrinsic MRE using naturally occurring cardiac pulsations, has shown stiffness values far below those obtained from post-mortem testing or externally actuated MRE. Such extreme softness allows a balance of elastic and inertial forces at even low frequencies, restoring uniqueness in viscoelastic iMRE inversion. Here, we demonstrate that viscoelastic iMRE can provide unique and stable stiffness estimates across frequencies, enabled by the brain’s ultra-soft nature.

**Method:** iMRE data was obtained for 8 healthy subjects from a previous 7T MRI study, and MRE data was obtained at 50 Hz for 38 healthy subjects from two prior studies. The elastic-to-inertial force ratio was calculated for all subjects and compared between intrinsic and extrinsic datasets. The convergence of the viscoelastic iMRE was evaluated across a range of initial conditions and was examined at approximately 1, 2, and 3 Hz.

**Results:** At a brain stiffness of 2600 Pa, the iMRE force ratio exceeded the MRE value by nearly four orders of magnitude, whereas at 4-35 Pa it fell to within approximately 0.5-2.5 orders of magnitude. Convergence was consistent across initial stiffness estimates. The mean storage modulus across all subjects was 5.29±0.95 Pa at 1 Hz, 34±17 Pa at 2 Hz, and 160±98 Pa at 3 Hz, consistent with previously reported frequency dependency.

**Conclusion:** Balanced force ratios, consistent convergence, and physiologically plausible results support uniqueness of the viscoelastic inversion. These findings resolve a key limitation in iMRE modeling and provide further evidence for the brain’s ultra-soft nature at low frequencies.

## 1. Introduction

Brain tissue is among the most complex tissues in the human body and is notoriously difficult to characterize. Emerging evidence suggests that the brain is exceptionally soft, and reported stiffness values have decreased progressively as measurement techniques have improved. Mechanical testing often yields values approximately in the range of 200 Pa to 1 kPa, positioning the brain as one of the softest organs in the body (Budday et al., 2020). However, such measurements are typically performed on post-mortem tissue, where the brain is removed from its natural, vascularized, and pressurized environment, potentially altering its mechanical properties. Magnetic resonance elastography (MRE) enables non-invasive *in vivo* assessment of brain stiffness, but standard externally actuated MRE often reports significantly higher values (2-3 kPa (Hiscox et al., 2020; Johnson et al., 2013)), as the measured stiffness increases strongly with actuation frequency (Herthum et al., 2021). Consequently, the natural softness of brain tissue in its normal environment and conditions remains difficult to establish using existing methodologies.

The feasibility of an adaptation of the MRE technique, called intrinsic MRE (iMRE), was recently shown, where the need for external actuation is circumvented by leveraging naturally occurring brain pulsations driven by the cardiac pulse (Burman Ingeberg et al., 2023; Solamen et al., 2021; Weaver et al., 2012; Zorgani et al., 2015). An advantage of iMRE is the ability to characterize brain tissue stiffness in a more physiologically natural state, rather than one altered by externally applied vibrations, and may thus provide more accurate measurements of the brain’s natural stiffness. However, when brain tissue is modeled as a viscoelastic material, as is common in standard MRE, the inversion problem at the low frequencies of cardiac pulsation becomes non-unique (M. McGarry et al., 2019). Here, the term non-unique refers to the fact that any scalar multiple of the viscoelastic properties provides indistinguishable tissue displacements, making it impossible to determine the tissue’s mechanical properties in an absolute sense. This ambiguity can lead to an inability for iMRE to detect clinical conditions leading to bulk stiffness changes, such as neurodegeneration or regeneration (Guo et al., 2019; McIlvain, Schneider, et al., 2022; Pavuluri et al., 2024; Takamura et al., 2020). The issue of non-uniqueness arises because, under high-stiffness and low-frequency conditions, elastic forces dominate over inertial forces, making the equilibrium conditions essentially quasi-static and valid for any constant scalar of the elasticity greater than unity (M. McGarry et al., 2019).

Recently, poroelastic inversion using iMRE was successfully performed. iMRE with poroelastic inversion preserves uniqueness due to fluid-matrix interaction (M. McGarry et al., 2019) and enables voxel-wise reconstruction of both hydraulic permeability (Burman Ingeberg et al., 2025). By contrast, iMRE implementations that impose a global hydraulic permeability scale the entire stiffness distribution (Burman Ingeberg et al., 2025), making this voxel-wise approach an important development that removes dependence on assumed global parameters. Using this approach, brain tissue stiffness was estimated at 5-6 Pa. Similarly, wavelength analysis with intrinsic activation steady-state MRE yielded an absolute complex shear modulus of 21 Pa (Herthum et al., 2021). Together, these findings indicate that the brain in its natural state is considerably softer than previously believed.

This insight carries important implications for computational modeling, as accurate simulations of brain mechanics rely on accurate stiffness estimates to reliably predict deformation under physiological and pathological conditions. It also highlights the brain’s vulnerability to even small mechanical stresses and the important role of fluid-structure interactions in normal brain function. Importantly, the brain’s extreme softness suggests that during the low-frequency cardiac vibrations measured in iMRE, elastic forces may no longer dominate inertial forces, potentially restoring uniqueness in the viscoelastic iMRE inversion process. Previous reports of non-uniqueness in viscoelastic modeling were likely influenced by the assumption that brain tissue exhibits much higher stiffness, i.e., values around 3 kPa typically observed in MRE (M. McGarry et al., 2019). In these cases, lower bounds on stiffness - typically around 100 Pa - were imposed to prevent the inversion from producing values considered unrealistically low (Burman Ingeberg et al., 2023; M. McGarry et al., 2019). However, these bounds can constrain convergence during inversion and artificially scale the resulting stiffness estimates, masking the underlying uniqueness of the solution.

In this study, we build on the observation that the brain is ultra-soft in its natural, unperturbed state. We demonstrate that, under these conditions, the balance between inertial and elastic forces in iMRE resembles that of conventional, externally actuated MRE. We further show that viscoelastic inversion in iMRE yields unique and stable in vivo property estimates without the need to impose a lower stiffness bound. Finally, we show that the viscoelastic model produces physically meaningful solutions, consistent with the frequency dependence observed by Herthum et al. (2021) by recovering property estimates at approximately 1, 2 and 3 Hz (using the first, second and third harmonic of the cardiac waveform). This ability to obtain unique iMRE solutions at low frequencies provides additional evidence of the brain’s extreme softness, with stiffness values low enough that the elastic forces of intrinsic, cardiac-driven brain deformation are balanced by inertial forces caused by the concomitant tissue motion.

## 2. Method

### 2.1 Intrinsic displacement data acquisition

The iMRE displacement data was previously obtained for a separate study by Adams et al., (2020). A brief overview of the acquisition method is provided here as full details are available in the original publication. All subjects gave written-informed consent for participation in the study, which was approved by the ethical review board of our institution.

Eight healthy young adults (three females; mean age: 27 ± 6 years) underwent imaging on a 7T MR scanner (Philips Healthcare) using a Displacement Encoding with Stimulated Echoes (DENSE) (Aletras et al., 1999) sequence with a 3D EPI readout. Cardiac synchronization was achieved through retrospective gating, using pulse oximetry to track the cardiac cycle. The spatial resolution of the acquired images was 1.95 mm × 1.95 mm × 2.2 mm (anterior-posterior, feet-head, right-left), and 20 temporal phases were reconstructed per cardiac cycle. Three separate acquisitions were conducted with displacement encoding (DENC) sensitivities of 0.175 mm, 0.175 mm, and 0.35 mm in the anterior-posterior (AP), right-left (RL), and foot-head (FH) directions, respectively. The time of acquisition per motion encoding direction was 144 heartbeats (corresponding to 2:24 minutes for an average heart rate of 60 beats per minute).

A Fast Fourier Transform (FFT) was applied to the temporal phases at each voxel to convert the displacement data into the frequency domain. Motion components at the first three harmonic frequencies, approximately 1 Hz, 2 Hz, and 3 Hz, were selected for further analysis. Although exact frequencies varied slightly depending on the subject’s resting hear rate, the terms “1 Hz”, “2 Hz” and “3 Hz” are used for simplicity and clarity.

### 2.2 Extrinsic displacement data acquisition

The externally actuated MRE displacement data (here referred to as extrinsic data) were obtained in separate, previously published studies (Caban-Rivera et al., 2024; Smith et al., 2022). A brief summary of the acquisition protocol is provided below as the full details can be found in the original publications. All participants provided informed, written consent to participate in this study approved by the Delaware Institutional Review Board.

A total of 38 subjects (19 males, 19 females; mean age: 46 ± 22 years) were scanned on a Siemens 3T Prisma MRI scanner. A pneumatic system (Resoundant, Inc., Rochester, MN) was used to introduce mechanical shear waves into the brain at 50 Hz. Two vibration sources were applied separately: a pillow driver producing AP motion, and a custom-designed LR actuator. The displacement fields from both actuations were acquired with a 3D multiband, multishot spiral sequence (McIlvain, Cerjanic, et al., 2022). Imaging parameters included 4 phase offsets, 2.0 mm isotropic voxel resolution, 240×240 mm^2^ FOV, 64 slices, and TR/TE = 2240/76 ms.

### 2.3 Ratio of elastic and inertial forces

The equation of motion for a heterogeneous viscoelastic material, without internal body forces, can be written in the frequency domain as

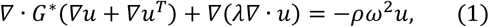

where *u* is the 3D complex-valued motion amplitude, *λ* is Lamé’s first constant [Pa], *ρ* is the density [kg/m^3^], and ω is the actuation frequency [rad/sec]. The complex-valued shear modulus *G*^∗^ [Pa] consists of the storage modulus *G*′ and the loss modulus *G*″, such that *G*^∗^ = *G*′ + *iG*″. The left-hand side of Equation (1) describes the elastic forces while the right-hand side describes the inertial forces.

Using this formulation, the ratio of elastic to inertial forces - capturing their balance - was calculated for three specific cases:

1. Extrinsic data at 2600 Pa: Extrinsic motion fields were evaluated at a global brain stiffness of 2600 Pa with an actuation frequency of 50 Hz. This value corresponds approximately to the average brain stiffness measured by standard MRE at 50 Hz (Hiscox et al., 2020)
2. Intrinsic data at 4 - 35 Pa: Intrinsic motion fields were evaluated across a range of global brain stiffnesses from 4 Pa to 35 Pa, representing the approximate lower and upper bounds obtained using intrinsic-actuation based methods at approximately 1 Hz (Burman Ingeberg et al., 2025; Herthum et al., 2021).
3. Intrinsic data at 2600 Pa: Intrinsic motion fields were also evaluated at a global brain stiffness of 2600 Pa with an actuation frequency of 1 Hz, reflecting previous assumptions on brain stiffness and allowing direct comparison to the extrinsic case.

For all cases, the density was set to *ρ* = 1000 kg m^−3^ and Lamé’s first constant was set to *λ* = 10*G*′ model near-incompressibility of soft tissue. The damping ratio *ξ* = *G*″/2*G*′ was set to 0.2, consistent with reported values for both extrinsic and intrinsic MRE (Burman Ingeberg et al., 2023; Hiscox et al., 2020).

### 2.4 Viscoelastic property estimation

For all following iMRE experiments, viscoelastic property values were estimated using a subzone-based non-linear inversion (NLI) scheme (M. D. J. McGarry et al., 2012; Van Houten et al., 2001), which incorporated quadratic hexahedral finite element models generated from each subject’s displacement data. A mesh size of 2.0 mm^3^ was used, along with a subzone size of 20 mm^3^ and 800 global iterations. A more detailed description of this methodology is provided by Burman Ingeberg et al., (2023). Viscoelastic inversion recovers the complex shear modulus *G*^∗^ = *G*′ + *iG*″.

### 2.5 Initial condition assessment

To assess convergence of the viscoelastic iMRE inversion at 1 Hz, five separate inversions were performed for each subject using different initial conditions for the storage and loss moduli (*G*′, *G*″): [1, 0.2], [5, 1], [10, 2], [20, 4], and [50, 10]. Under conditions where the inverse problem is well-posed and the solution is unique; converged property estimates should be independent of the initial condition. No lower bounds were imposed on the storage and loss moduli in order to avoid artificially constraining the inversion or biasing the solution toward a predefined minimum.

### 2.6 Viscoelastic frequency dependence

The frequency dependence of the viscoelastic model was assessed by performing viscoelastic inversion on the 1 Hz, 2 Hz and 3 Hz motion components for each subject. If the model yields unique and physically meaningful solutions, the recovered property values are expected to increase with frequency (Herthum et al., 2021). Analysis was limited to the first three harmonic components due to a substantial decline in signal-to-noise ratio (SNR) at higher frequencies. For all inversions, the initial conditions for *G*′and *G*″ were set to 30 Pa and 6 Pa, respectively. Lamé’s first constant was set 10 times the initial guess for *G*′.

The resulting mean global property estimates were directly compared to a dispersion curve measured by Herthum et al., (2021) by converting the estimated complex shear modulus to shear wave speed (SWS) using

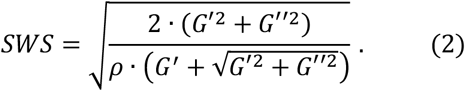

Here, *ρ* represents the tissue density and was set to *ρ* = 1000 kg m^−3^. Following the approach of Herthum et al., (2021) both viscous- and Kelvin-Voigt models were fitted to our data and to Herthum’s data using a nonlinear least-squares algorithm. The viscous model can be expressed as

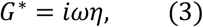

and the Kelvin-Voigt model by

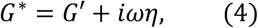

where *η* is the viscosity. For the least-squares fitting, data above 3 Hz were obtained from Herthum et al. (2021), data in the 2-3 Hz range were taken from the iMRE in the present study, and the 1 Hz data was calculated as the mean of both datasets.

## 3 Results

### 3.1 Balance of inertial and elastic forces

The ratio between the inertial and elastic forces was successfully calculated using Equation 1 for all subjects for both intrinsic and extrinsic displacement data. The mean ratio (inertial/elastic) across all subjects was 107 ± 34 for extrinsic actuation at 2600 Pa, spanned from 627 to 5490 for intrinsic actuation at 4 - 35 Pa, and 4.1 × 10^5^ ± 4.7 × 10^4^ for intrinsic actuation at 2600 Pa. Figure 1 presents boxplots displaying the distribution of force ratios across subjects for each respective situation. Intrinsic actuation within the 4-35 Pa stiffness range yielded force ratios at the lowest stiffness that were within half an order of magnitude of those observed for extrinsic actuation at 2600 Pa. At the upper end of the range, the force ratios increased to approximately 2.5 orders of magnitude. In contrast, intrinsic actuation at 2600 Pa produced ratios nearly four orders of magnitude higher than extrinsic actuation at the same stiffness.

**Figure 1:**
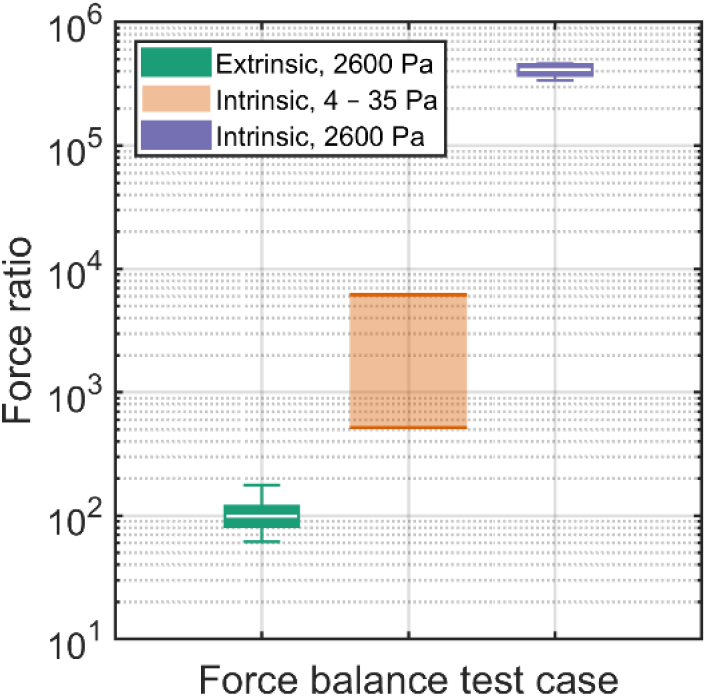
Boxplots of the distribution of force ratios across subjects for three cases: extrinsic actuation at 2600 Pa and 50 Hz, intrinsic actuation at 1 Hz with global brain stiffness spanning 4-35 Pa (visualized as a shaded range in the figure), and intrinsic actuation at 1 Hz at a brain stiffness of 2600 Pa. In each boxplot, the edges represent the 25th and 75th percentiles, the whiskers show the minimum and maximum values within 1.5x the interquartile range, and the white line indicates the mean.

**Figure 2:**
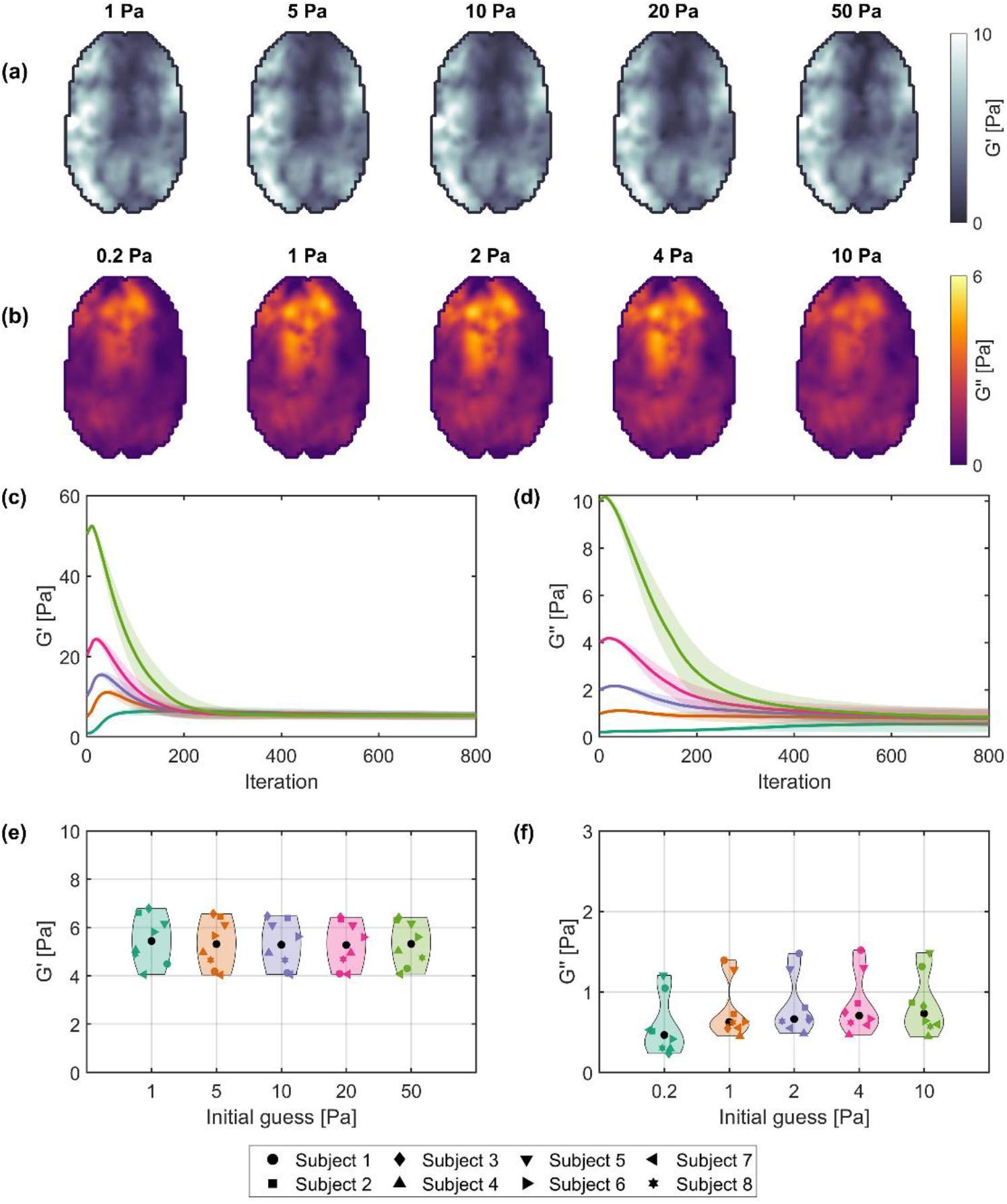
Property maps for Subject 1 at different initial conditions: (a) storage modulus and (b) loss modulus. The loss modulus failed to fully stabilize when initialized at 0.2 Pa due to slower convergence. Convergence plots across all subjects for the storage modulus (c) and loss modulus (d) across initial conditions. Solid lines indicate mean values across subjects, and shaded areas represent the standard deviation. Violin plots in (e) and (f) show the distribution of mean storage and loss moduli, respectively, at the final iteration.

### 3.2 Initial condition assessment

The iMRE viscoelastic properties were successfully reconstructed for all subjects and across all initial conditions. The mean global storage and loss moduli across subjects are presented in Table 1. Figure 1a shows storage modulus maps, and Figure 1b shows loss modulus maps for a representative subject under different initial conditions. Convergence plots for both moduli are presented in Figures 1c and 1d, while Figures 1e and 1f display violin plots summarizing the mean values across subjects for the storage and loss moduli, respectively. Regardless of the initial conditions, all subjects converged toward similar mean property values. However, the loss modulus converged considerably slower than the storage modulus, failing to fully stabilize after 800 iterations when initialized at particularly low values (e.g., 0.2 Pa). These effects are evident in both the loss modulus maps and the decreased values observed at the 800th iteration for the 0.2 Pa initialization. Consequently, the storage modulus exhibited slightly higher final values for the 1 Pa initial condition. Despite these differences, the storage modulus retained a consistent spatial distribution across different initial conditions.

**Table 1:** Global storage and loss moduli (mean ± standard deviation across subjects) for different initial conditions.

| Initial conditions<br>[$G'$, $G''$] [Pa] | [1, 0.2] | [5, 1] | [10, 2] | [20, 4] | [50, 10] |
| --- | --- | --- | --- | --- | --- |
| $G'$ [Pa] | 5.48 ± 0.94 | 5.33 ± 0.94 | 5.29 ± 0.92 | 5.28 ± 0.91 | 5.33 ± 0.86 |
| $G''$ [Pa] | 0.57 ± 0.34 | 0.78 ± 0.33 | 0.82 ± 0.34 | 0.85 ± 0.35 | 0.84 ± 0.35 |

### 3.3 Viscoelastic frequency dependence

The storage and loss moduli were successfully reconstructed for the 1 Hz, 2 Hz, and 3 Hz motion components for all subjects. Representative axial slices of the corresponding storage and loss modulus maps are provided in Figures S1 and S2 of the supplementary material. The mean storage modulus across all subjects after 800 iterations was 5.29 ± 0.95 Pa at 1 Hz, 34 ± 17 Pa at 2 Hz, and 160 ± 98 Pa at 3 Hz. Similarly, the mean loss modulus across all subjects was 0.85 ± 0.38 Pa at 1 Hz, 6.8 ± 3.8 Pa at 2 Hz, and 12.8 ± 7.6 Pa at 3 Hz. Figure 3 shows the corresponding mean convergence plots. Both the storage and loss moduli exhibited an increase with rising frequency, consistent with frequency dependent behavior of viscoelastic material. The variability, as reflected by the standard deviation across subjects, increased considerably at higher frequencies, which can be attributed to the progressive reduction in signal-to-noise ratio (SNR) with each incremental frequency.

**Figure 3:**
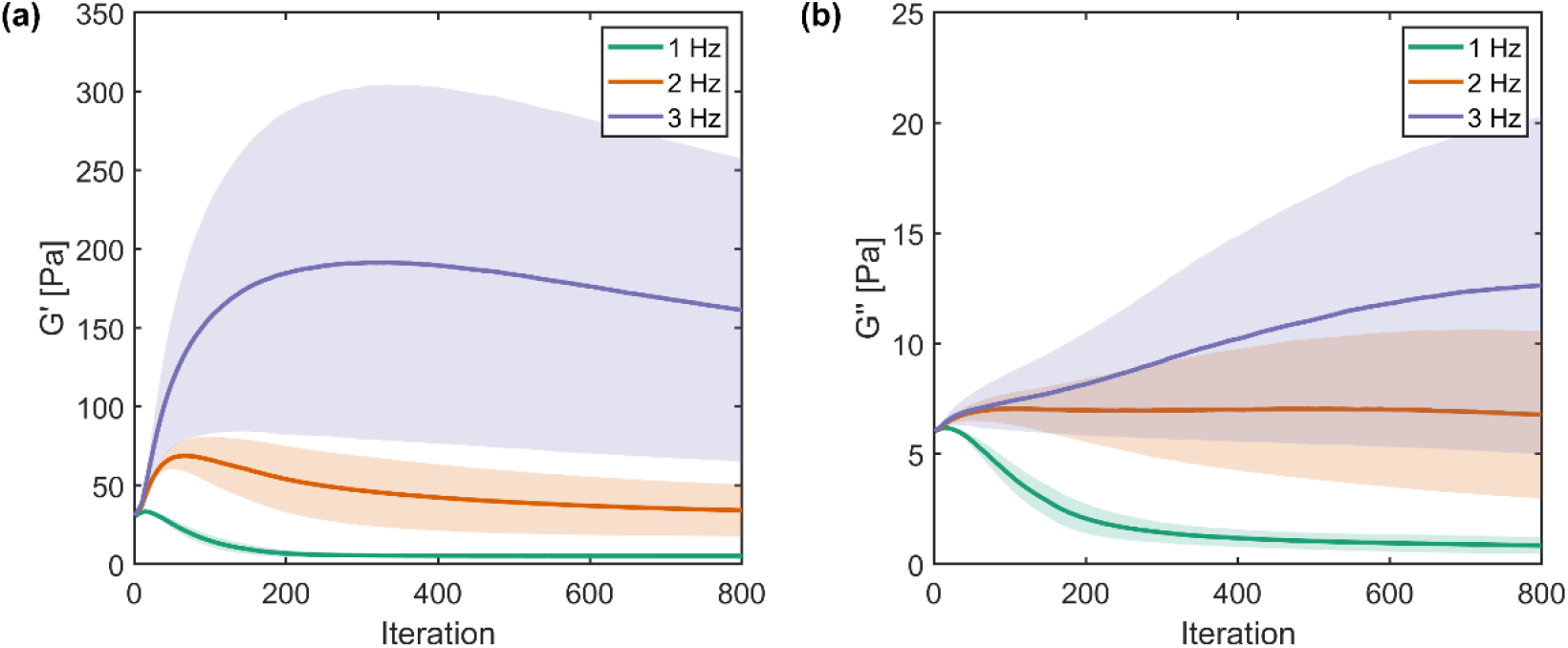
Convergence plots across for the storage modulus (a) and loss modulus (b) for the 1 Hz, 2 Hz, and 3 Hz motion components. The solid line represents the mean across all subjects, with the shaded area representing the corresponding standard deviation.

Both the viscous and Kelvin-Voigt models were fitted successfully, resulting in *η* = 6.1 Pa ⋅ s for the viscous model and *G*′ = 40 Pa, *η* = 6.5 Pa ⋅ s for the Kelvin-Voigt model. The corresponding model fits are presented in Figure 4, together with shear wave speeds calculated from the storage and loss moduli using Equation 2, and higher-frequency measurements reported by Herthum et al., (2021).

**Figure 4:**
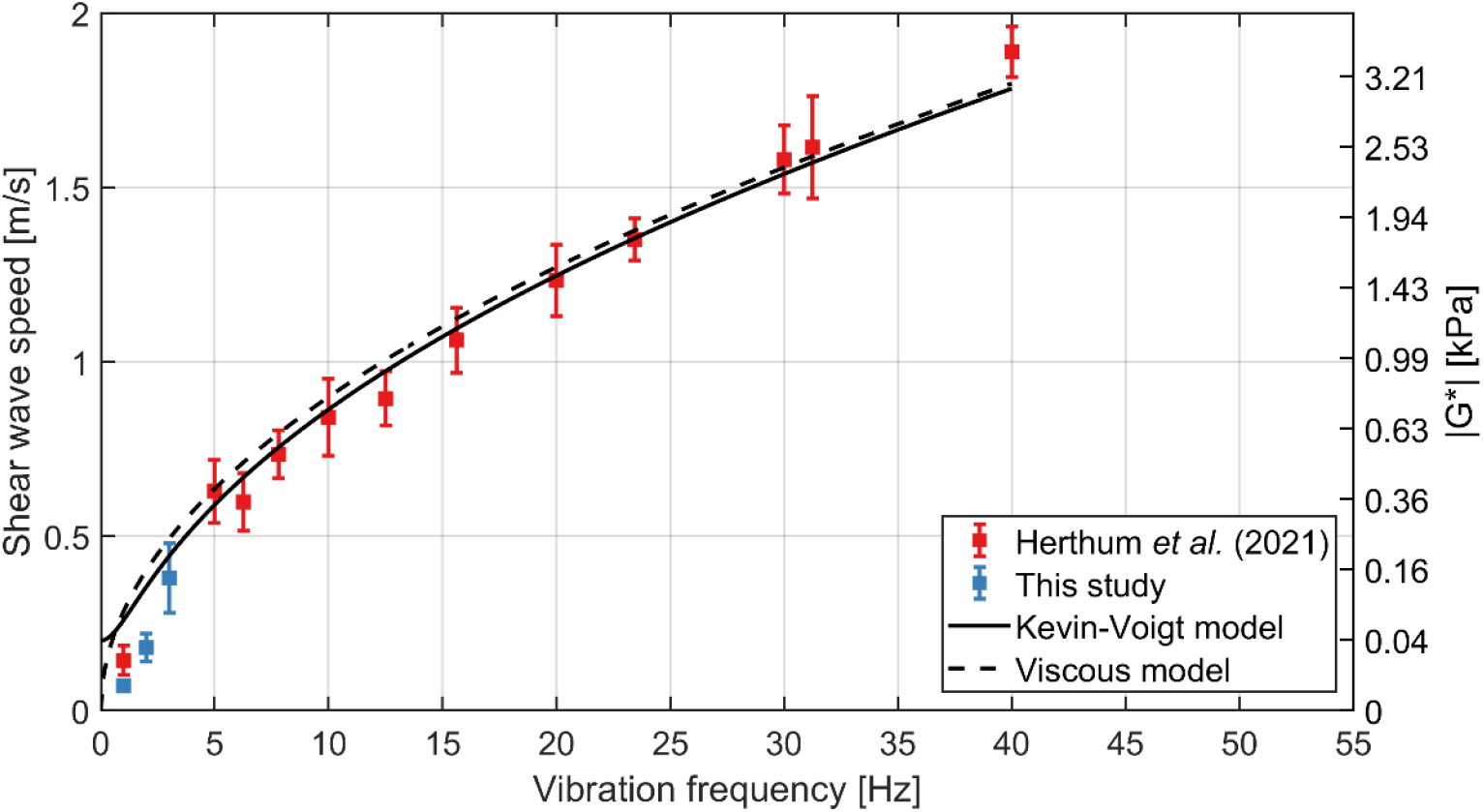
Dispersion curve showing the frequency dependence of shear wave speed (left y-axis) or absolute complex shear modulus |***G***^∗^| (right y-axis) for the present study and for the data reported by Herthum et al., (2021). The dots indicate mean values while whiskers indicate standard deviation. Solid and dashed lines show model fits using a Kelvin–Voigt and a viscous model, respectively. The right y-axis was calculated using 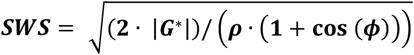 with ***ϕ*** = **0** to provide an approximate conversion to the magnitude of the complex shear modulus.

## 4 Discussion

Viscoelastic inversion in low-frequency settings such as iMRE has been considered ill-posed due to an assumed imbalance between elastic and inertial forces, leading to non-unique solutions. To address this, inversion pipelines commonly impose lower-bound stiffness constraints to enforce convergence. However, recent hints of the ultra-soft nature of the brain in its natural state, and the findings of our current study challenge the necessity of this assumption. We present multiple lines of evidence demonstrating that viscoelastic inversion at 1 Hz yields unique and physiologically meaningful stiffness estimates, without the need for artificial regularization via stiffness floors.

The uniqueness of viscoelastic solutions in iMRE depends on the balance between elastic and inertial forces, which in turn is influenced by tissue stiffness and actuation frequency. In this analysis, global stiffness values spanning 4 - 35 Pa were used to compute the elastic-inertial force balance, representing literature-based estimates for brain stiffness under intrinsic actuation without imposing arbitrary lower boundaries (Burman Ingeberg et al., 2025; Herthum et al., 2021). The resulting ratios ranged from values within approximately half an order of magnitude of those observed for extrinsic actuation at the lowest stiffness, to up to 2.5 orders of magnitude higher at the upper end of the range. In contrast, assuming a stiffness of 2600 Pa for intrinsic actuation yielded a force ratio more than three orders of magnitude greater than in the extrinsic case, placing the system deep within an elastic-dominated regime where non-uniqueness is expected. These results suggest that intrinsic actuation at low stiffnesses produces force balances more comparable to extrinsic conditions, whereas increasing stiffness markedly amplifies the imbalance and likely contributes to non-unique viscoelastic reconstructions. This may help explain why previous studies have reported non-uniqueness in viscoelastic inversions at 1 Hz, where assumed stiffness values were as high as 3.3 kPa (M. McGarry et al., 2019).

If the viscoelastic model yields unique solutions, the inversion process should converge to similar property maps regardless of the initial conditions, provided sufficient time for convergence. In this study we observed that both the storage and loss moduli reached nearly identical global mean values across a range of different initializations. This led to nearly identical global means and spatial distributions of the storage and loss moduli, regardless of initial conditions, even without stiffness lower bounds that could artificially enforce consistency. Furthermore, very similar mean stiffness values were obtained across subjects, with the viscoelastic inversion yielding a mean of 5.29 Pa, closely matching previously obtained poroelastic (6.09 Pa) and poroviscoelastic (5.33 Pa) inversions (Burman Ingeberg et al., 2025). This also highlights the tendency for slightly higher stiffness estimates when the loss modulus is not modeled, as in the purely poroelastic inversion.

However, when the loss modulus was initialized at 0.2 Pa, the inversion did not have sufficient time to fully converge, leading to noticeable differences in the final results. In general, the loss modulus converged considerably more slowly than the storage modulus, contributing to larger discrepancies in the property maps at the final iteration. This behavior is likely related to the use of a purely viscoelastic model at low actuation frequencies, where poroelastic effects become more prominent (M. McGarry et al., 2019). In such conditions, poroelastic damping in the brain is inadequately captured by a viscous model, causing the loss modulus to poorly represent the observed attenuation and consequently to converge more slowly. This also likely explains the slight increase observed in the global mean storage modulus for the 1 Pa initial condition, highlighting the importance of appropriate initial conditions for effective convergence, particularly for the loss modulus.

The viscoelastic properties of brain tissue generally show a strong frequency dependence. If the viscoelastic model yields unique, physically meaningful solutions, an increase in property values with increasing frequency would be expected. This trend was consistently observed across all subjects, providing additional support for uniqueness. However, the relative amplitude of the motion decreases for higher harmonics, leading to a corresponding reduction in the signal-to-noise ratio (SNR) which considerably affects the quality of the property estimates. This, in turn, can make convergence difficult and was reflected in the larger inter-subject variability observed in the storage and loss moduli at 2 Hz and 3 Hz. This effect was most pronounced at 3 Hz, where the final property estimates should be interpreted with caution.

The observed frequency dependence closely resembled the dispersion behavior reported by Herthum et al., (2021), as shown in Figure 4, with a steep increase in stiffness at low frequencies. The measured values at the lowest frequencies were, however, consistently lower than predicted by the both the viscous- and Kelvin-Voigt models. This deviation is expected, as the viscoelastic model provides an incomplete description of brain mechanics at low actuation frequencies, where poroelastic effects start to dominate. When the fluid equilibration timescale approaches the motion period, as at 1 Hz, poroelastic modeling is required for accurate reconstruction. At higher frequencies, the equilibration time becomes much longer than the oscillation period, and the viscoelastic model serves as an adequate and simpler approximation of poroelastic behavior (M. McGarry et al., 2019; M. D. J. McGarry et al., 2015).

The evidence presented above strongly supports the ability of viscoelastic inversion to yield unique solutions in iMRE. Notably, the very existence of such unique solutions at low frequencies can itself be interpreted as indirect evidence of the brain’s ultra-soft nature, as only in this regime do elastic and inertial forces remain sufficiently balanced to prevent non-uniqueness. To better understand this regime, it is useful to compare iMRE-derived stiffness values with those obtained from mechanical testing, which are typically much higher (0.4–1.4 kPa (Budday et al., 2017)). Several factors likely contribute to this discrepancy. First, mechanical testing is generally performed on post-mortem brain tissue. While stiffness has been reported to remain relatively stable for hours to days post-mortem under appropriate preservation (Budday et al., 2017, 2020), other studies using nanoindentation in bovine tissue observed substantial early stiffening, with increases of 26 % in the cerebrum and 58 % in the corpus callosum after only 3 minutes, rising to 41 % and 142 % after 45 minutes, respectively (Weickenmeier et al., 2018). Second, mechanical testing and iMRE probably probe different stress regimes. Whereas mechanical testing typically applies stresses on the order of 0.1 – 2 kPa (Budday et al., 2020; Chatelin et al., 2010; Hou et al., 2025), iMRE relies on subtle cardiac-induced brain pulsations. Although precise estimates of such stresses have yet to be determined, they are likely considerably smaller than that of mechanical testing, producing strains of 10^−4^ - 10^−2^ (Adams et al., 2020; Hirsch et al., 2013; Sloots et al., 2021), compared to up to 10% used in mechanical testing (Budday et al., 2020). Because biological tissue generally does not exhibit a linear stress-strain relationship over wider stress ranges, lower applied stresses may correspond to lower apparent stiffnesses. Third, the brain is a highly porous solid matrix saturated with interstitial and vascular fluids. At the low actuation frequencies used in iMRE, these fluids have sufficient time to redistribute in response to deformation, effectively reducing the tissue’s mechanical resistance. Indeed, recent work has shown that recovery of brain tissue after indentation occurs on the order of seconds (Liu et al., 2025) supporting the view that lower actuation frequencies naturally capture this redistribution and explaining why porous models are most appropriate for interpreting measurements in this regime (M. McGarry et al., 2019).

While the points discussed above may account for some of the discrepancies in stiffness estimates between iMRE and mechanical testing, they are unlikely to provide a complete explanation. Rather, iMRE appears to capture a distinct aspect of brain tissue mechanics, where the brain’s suspension and perfusion by fluid in a living, dynamic system play a major role, producing consistently low stiffness values that suggest an almost fluid-like response at low frequencies. Recent work on poroviscoelastic modeling highlights the importance of Lamé’s first constant, representing resistance to volumetric deformation, in governing fluid flow (Greiner et al., 2024). Higher values, corresponding to a stiffer brain, restrict fluid movement and lead to physically unrealistic behavior. This suggests that low stiffness may be key in maintaining healthy fluid dynamics, both in waste clearance and in vascular perfusion. In this line, one could speculate that lower brain stiffness is a means of optimizing fluid regulation. Taken together, these observations indicate that the brain’s mechanical and structural integrity depends on both the surrounding fluid and the protective enclosure provided by the skull. Unlike organs such as the liver, which lack rigid external protection, the skull allows the brain to maintain its exceptionally low stiffness without compromising structural stability.

An intriguing consequence of this perspective is that systemic factors, such as cardiovascular dynamics, could modulate effective brain stiffness. For instance, lower resting heart rates - particularly during sleep - may effectively soften the brain. This, in turn, could further facilitate fluid movement and waste clearance, consistent with observations that metabolic waste removal in the brain is enhanced during sleep (Xie et al., 2013). Recent studies further support this link, demonstrating that lower heart rates are associated with enhanced clearance (Dagum et al., 2025; Hablitz et al., 2019). However, these notions remain speculative, and further work is needed to clarify the precise relationships between brain stiffness, cardiovascular dynamics, and fluid clearance.

While currently viscoelastic, poroelastic and poroviscoelastic inversions all converge towards similar stiffness estimates of approximately 5-6 Pa, they differ considerably to the 21 Pa measured by Herthum et al., (2021). However, their estimation is based on a purely elastic model, which ignores viscous effects and may produce higher apparent stiffnesses. This phenomenon would be consistent with our earlier observation that the purely poroelastic model estimates higher stiffness than the viscoelastic and poroviscoelastic model. That said, it is also possible that our own estimates slightly underestimate the true values. Therefore, a limitation of this study is that our current iMRE framework has not yet been validated in phantom experiments. Therefore, future work aims to perform validation studies in sufficiently soft, well-characterized phantoms where the elastic-inertial force balance permits unique viscoelastic reconstructions.

## 5 Conclusion

This study demonstrates that viscoelastic inversion in intrinsic MR elastography can yield unique and physiologically meaningful property estimates. Due to the ultra-soft nature of the brain under low actuation frequencies, we show that the balance between elastic and inertial forces more closely resembles that of conventional MRE, thereby restoring uniqueness in the inversion process. The resulting viscoelastic properties converge consistently across varying initial conditions, display frequency dependence consistent with prior literature, and provide similar stiffness estimates as poroelastic and poroviscoelastic models - all without imposing artificial stiffness bounds. This work further establishes the brain as ultra-soft in its intact state under low actuation frequencies and addresses a major limitation of the viscoelastic model in intrinsic MR elastography.

## 6 Acknowledgements

We would like to thank Dr. Curtis Johnson for kindly providing the dataset that supported this research, and to Dr. Helge Herthum for contributing the data used in Figure 4.

This work was supported by a Vici Grant from the Netherlands Organization for Scientific Research (NWO) awarded to Jaco J.M. Zwanenburg under grant agreement no. 18674.

## Data availability statement

The data presented in this study are available from the corresponding author upon request.

## Ethics approval statement

The imaging performed for this study was done under the research proposal “MRI protocol development for all field strengths”, approved by the local Ethics Review Board of the UMC Utrecht (METC 15-466).

## Patient consent statement

All subjects gave written-informed consent for participation in this study, which was approved by the ethical review board of UMC Utrecht.

