## Supplementary Material for "Evidence for the ultra-soft brain provided by uniqueness in intrinsic MR elastography"

### Supplementary files

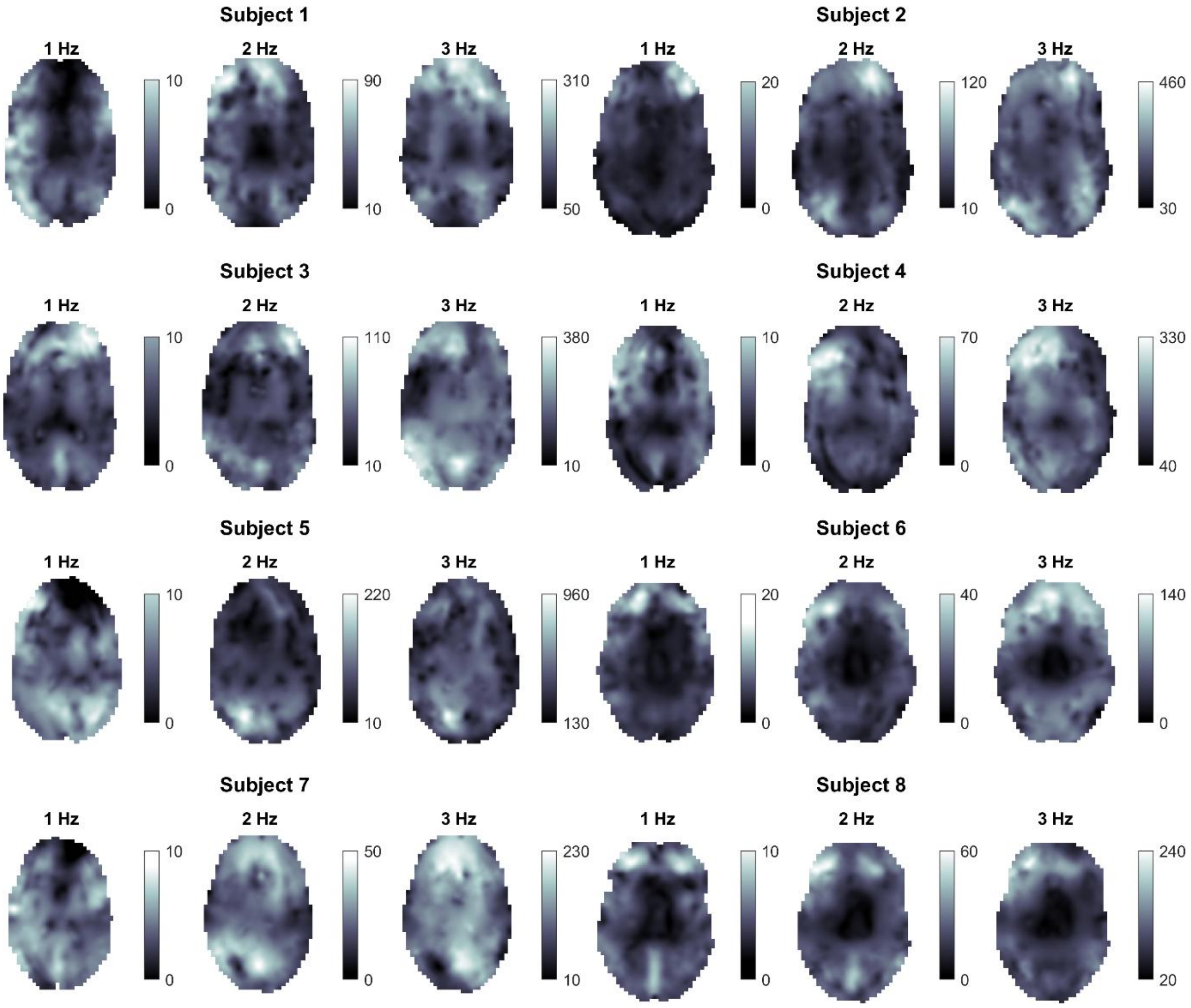


**Figure S1:** Representative axial slices of storage modulus maps at approximately 1, 2, and 3 Hz for all 8 subjects.

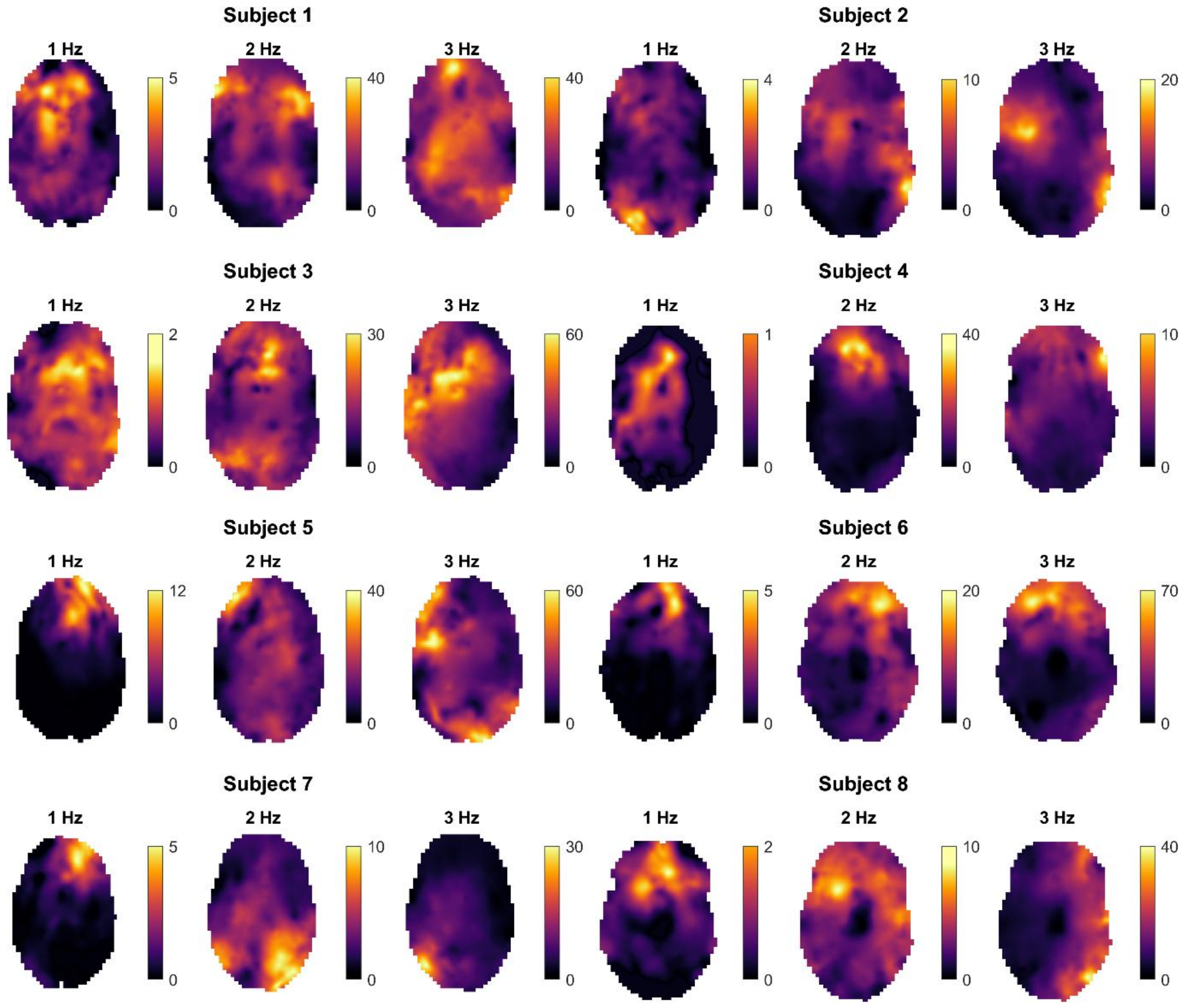


**Figure S2:** Representative axial slices of loss modulus maps at approximately 1, 2, and 3 Hz for all 8 subjects.
